# Algorithm for Reliable Detection of Beat Onsets in Cerebral Blood Flow Velocity Signals

**DOI:** 10.1101/364356

**Authors:** Nicolas Canac, Mina Ranjbaran, Michael J. O’Brien, Shadnaz Asgari, Fabien Scalzo, Samuel G. Thorpe, Kian Jalaleddini, Corey M. Thibeault, Seth J. Wilk, Robert B. Hamilton

## Abstract

Transcranial Doppler (TCD) ultrasound has been demonstrated to be a valuable tool for assessing cerebral hemodynamics via measurement of cerebral blood flow velocity (CBFV), with a number of established clinical indications. However, CBFV waveform analysis depends on reliable pulse onset detection, an inherently difficult task for CBFV signals acquired via TCD. We study the application of a new algorithm for CBFV pulse segmentation, which locates pulse onsets in a sequential manner using a moving difference filter and adaptive thresholding. The test data set used in this study consists of 92,012 annotated CBFV pulses, whose quality is representative of real world data. On this test set, the algorithm achieves a true positive rate of 99.998% (2 false negatives), positive predictive value of 99.998% (2 false positives), and mean temporal offset error of 6.10 ± 4.75 ms. We do note that in this context, the way in which true positives, false positives, and false negatives are defined caries some nuance, so care should be taken when drawing comparisons to other algorithms. Additionally, we find that 97.8% and 99.5% of onsets are detected within 10 ms and 30 ms, respectively, of the true onsets. The algorithm’s performance in spite of the large degree of variation in signal quality and waveform morphology present in the test data suggests that it may serve as a valuable tool for the accurate and reliable identification of CBFV pulse onsets in neurocritical care settings.

## I. Introduction

Assessment of cerebrovascular function is imperative in the diagnosis and management of numerous conditions common in neurological care. Transcranial Doppler (TCD) ultrasound has been used since the 1980s to measure cerebral blood flow velocity (CBFV), which can serve as a means of assessing of cerebral hemodynamics [1], [2]. TCD has established clinical indications for subarachnoid hemorrhage, cerebral vasospasm, acute ischemic stroke, stenosed or occluded intracranial vessels, and sickle cell disease [3], [4], [5]. In addition, monitoring of CBFV may be useful as a tool for noninvasive intracranial pressure monitoring and the assessment of mild traumatic brain injury [6], [7], [8], [9], [10].

CBFV waveform analysis often utilizes individual CBFV pulse morphology and thus depends heavily on reliable pulse onset detection. Accurate pulse delineation in CBFV waveforms presents a significant challenge for a number of reasons. There is an inherent difficulty in TCD measurement due to the scatter and attenuation of the signal by the skull, resulting in a relatively low signal to noise ratio [11], [12]. Additionally, TCD is highly operator dependent and relies on the operator’s ability to locate the acoustic window and insonate the appropriate vessel within a cerebrovasculature which may vary widely from patient to patient. Furthermore, CBFV signals are particularly prone to noise artifacts as a result of motion of the (not uncommonly handheld) probe or subject. Finally, the wide range of possible waveform morphologies that can occur in CBFV signals, particularly in pathological populations, presents yet another challenge, as it may be difficult to rely on the presence of any one uniform and consistent feature to aid in onset detection.

Despite these difficulties, TCD remains a compelling diagnostic tool due to being relatively inexpensive, noninvasive, fast, and portable. A solution to the problem of extracting individual CBFV pulses coupled with a reliable and accessible method of CBFV monitoring could open up numerous opportunities for improved neurocritical care. In this study, we focus on the former problem of accurate beat-to-beat segmentation. The problem can be reduced to locating the starting point of each beat or pulse, the so-called pulse onset, as we will refer to it, or pulse foot [13].

Though a number of methods have been developed for pulse onset detection for other physiological signals, including arterial blood pressure (ABP) and photoplethysmogram (PPG) [14], [15], [16], [17], [18], relatively few methods have been applied to CBFV signals. Those that do often require information from another complementary signal, such as electrocardiogram (ECG) [19]. The authors in [20] applied an adaptive thresholding strategy to CBFV pulses that required no other information and obtained a true positive rate (TPR) of 93.1% and a positive predictive value (PPV) of 93.3%.

In this paper, we present a new algorithm for pulse onset detection in CBFV waveforms that does not require any additional complementary signals and seeks to improve on the performance of previous methods. This algorithm incorporates a moving difference filter (MDF) and adaptive thresholding to indentify windows in which onsets are likely to occur. The algorithm then performs a local search within each window to determine the most probable position for the onset. Finally, the algorithm seeks to identify and correct errors by applying outlier detection strategies. The result of this method is a substantial improvement over existing pulse onset detection algorithms.

We begin by describing the data set that was used to test the performance of the algorithm. We then present a detailed description of the process of defining search windows, locating likely pulse onsets within those windows, and handling onset detection errors. We then move on to a presentation of the performance of the algorithm on the test data set, including an analysis of the impact of free parameters. Finally, we conclude by summarizing these results and offer some commentary on the potential use cases and robustness of the algorithm.

## II. Data set

The data set used in this work is identical to the one used in [20] and consists of 92,794 annotated CBFV pulses collected by a trained sonographer from subarachnoid hemorrhage patients admitted to the UCLA Medical Center. The subject group consisted of 108 patients (42 female), aged 30-64, with an average age of 48. However, due to unusable or unrecognizable CBFV signals resulting from extreme aliasing and other user errors, six total scans were excluded from this analysis, resulting in a total of 92,012 annotated pulses. The patients were consented under the protocol approved by the UCLA Internal Review Board (#10-001331) prior to analyzing any of their data. Electrocardiogram (ECG) signals were also collected, and, being a simpler waveform for pulse detection, were used for the purpose of facilitating the process of annotating pulse onsets. Onset locations were found using the R peak of the QRS complex in the ECG signal as discussed in [13] and [20]. The resulting annotations were then manually inspected for accuracy and corrected where necessary. These annotations are treated as the ground truth when evaluating the performance of the algorithm.

A potential feature of the data set is that it does not represent an idealized scenario, as a significant portion of the data would be considered relatively poor quality. This, coupled with a broad range of different waveform morphologies, provides a very robust test of the performance of the algorithm. In real world applications, poor quality data and high morphological variance are common due to the inherent challenges in TCD measurement outlined previously. As such, strong performance on perfect data is not necessarily informative as to how well an algorithm will perform in the field, so testing on data that realistically spans the spectrum of possible signals is imperative to properly evaluate performance. Fig. 1 illustrates a number of examples pulled directly from the test data set, which are meant to provide a representative survey of the range of signals encountered, both in terms of inherent signal quality and morphological variation.

**Fig. 1.**
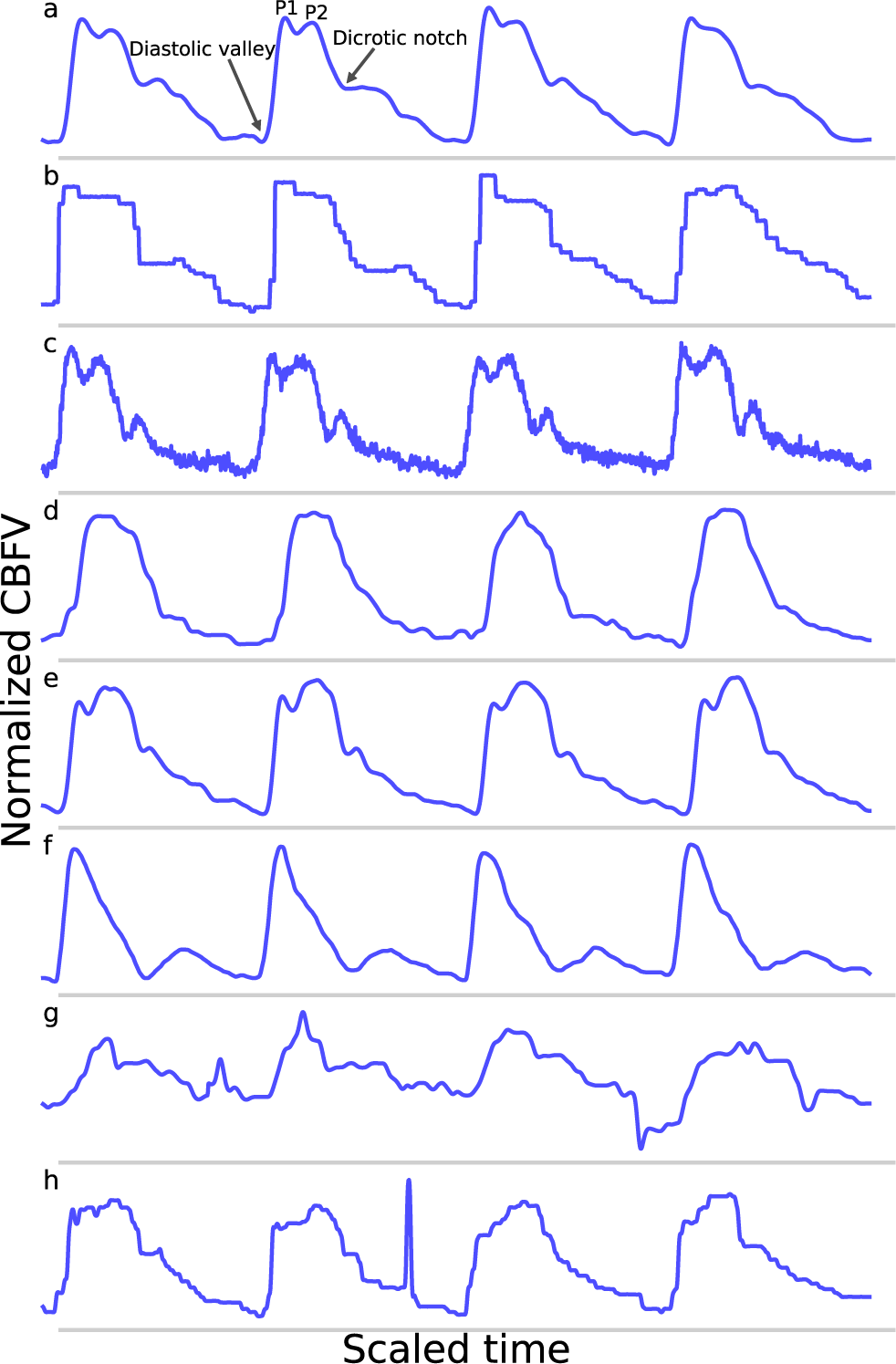
Shown are samples taken from the data set. Panel (a) shows a good quality, healthy waveform for reference. Panel (b) demonstrates a common measurement error that results in signal with a digitized appearance. Panel (c) shows a signal with a significant high frequency noise component. Panel (d) shows an example of a blunted waveform. Panel (e) shows a waveform with an elevated P2. Panel (f) shows a potentially stenotic waveform. Panel (g) demonstrates a signal with extremely poor quality. And finally, panel (h) illustrates a signal with a sharp noise spike.

For reference, Fig. 1(a) shows an example of a relatively good quality signal exhibiting what we would consider a normal, healthy waveform. Several common morphological landmarks have also been labeled for reference: the diastolic valley, dicrotic notch, first peak (P1), and second peak (P2). Fig. 1(b-c) illustrate two very common types of noise found in the data set. Fig. 1(b) may be the result of sampling error causing a digitized appearance, and Fig. 1(c) contains a significant high frequency noise component. Fig. 1(d-f) represents what one might consider to be “pathological” signal morphologies. While it is not clear that these signals represent a definitive underlying hemodynamic pathology, they nevertheless diverge in important ways from what would be considered a standard, healthy waveform and serve to demonstrate the importance of robustness to morphological variation for any onset detection algorithm. Fig. 1(d) shows a typical blunted waveform as described and classified in [21], in which independent P1 and P2 are not clearly identifiable. Fig. 1(e) shows another not uncommonly seen morphological variant in which the second peak (P2) is consistently higher than the first peak (P1). Fig. 1(f) depicts a signal where the dicrotic notch and diastolic valley of each beat are nearly equal, resulting in two clearly distinct peaks of different amplitudes, a characteristic that may be associated with stenotic waveforms defined in [21]. Fig. 1(g) is meant to be illustrative of the types of extremely poor quality signals encountered in the data, and Fig. 1(h) exhibits a signal with a sharp noise spike, another common feature that might trick an algorithm into incorrectly identifying the start of a new beat.

The example segments in Fig. 1 are meant to be illustrative of the types of noise, signal morphologies, and quality that are present in the data set that inhibit accurate onset detection. However, these examples should not be taken to be fully representative of the entire data set. In general, the signals span a range, from relatively good signals with normal morphologies to very poor signals with pathological morphologies. This is one reason why this data set offers a very robust test of the performance of the algorithm. If the same algorithm can perform well for a wide range of signal qualities and signal morphologies, then we can be more confident that it may perform well in a real world or clinical setting where differences resulting from sonographers, patients, or equipment can all contribute to increased scan to scan variance.

## III. Method

The algorithm consists of six primary steps. First, the signal is pass-band filtered to remove noise. Then, a moving difference filter (MDF), defined in eq. (1), is applied to the signal. Afterwards, the MDF signal is used to help define search windows on the CBFV signal in which to look for candidate onset locations. The beat onsets are then located within these search windows. Once all candidate onsets have been identified in this manner, a beat length analysis step is performed to identify and handle possible errors. Finally, a beat alignment step is performed to fine tune the precise locations of the onsets. An overview of this general process is illustrated in Fig. 2. Each of these steps is described in more detail in the subsections below.

**Fig. 2.**
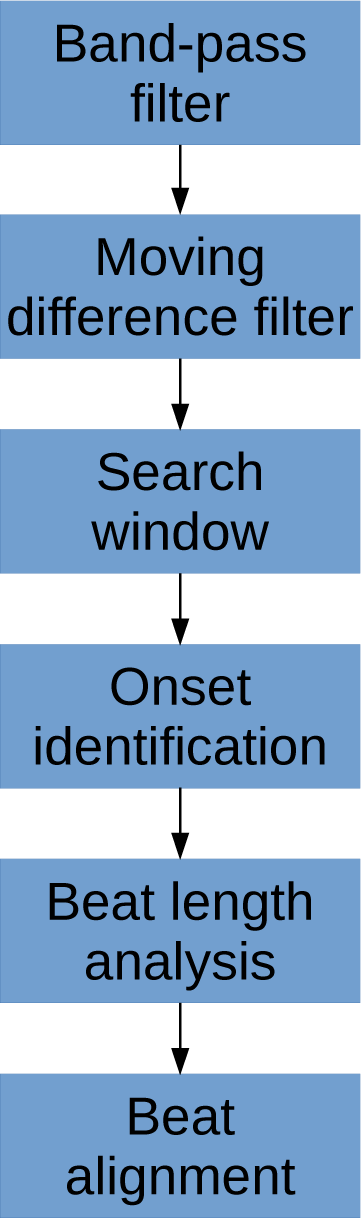
The flowchart showing a high level overview of the steps involved in identifying beat onsets.

### A. Band-pass filter

The signal is band-pass filtered using a fourth-order Butterworth filter with a passband range of 0.5-10 Hz. This filtering is done primarily to remove high frequency noise, which can complicate the matter of finding local extrema, as well as low frequency motion or breathing artifacts. We feel that this choice of passband range preserves all of the physiologically relevant features for onset detection. This filtered signal is passed along to the subsequent steps to be used for locating onsets.

### B. Moving difference filter

A moving difference filter (MDF) is applied to the signal in order to enhance the sharp upslope that defines the start of a typical CBFV pulse. The MDF is defined as follows:

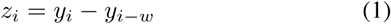

where *w* is the length of the analyzing window and *y* is the filtered CBFV signal. The MDF serves a similar purpose as the slope sum function used in [14], but we find that the MDF performs better in terms of filtering noise and preventing false detections. The size of the analyzing window should be roughly equal to the length of the initial upslope of a typical CBFV pulse in order to maximally enhance the upslope and be less sensistive to noise artifacts. In this work, the analyzing window is taken to be 150 ms, though the performance of the algorithm is fairly robust to slight changes around this value as discussed in more detail in Section IV-B. An example of a CBFV waveform with a detected pulse is shown in Fig. 3(a) with its corresponding MDF signal displayed in Fig. 3(b).

**Fig. 3.**
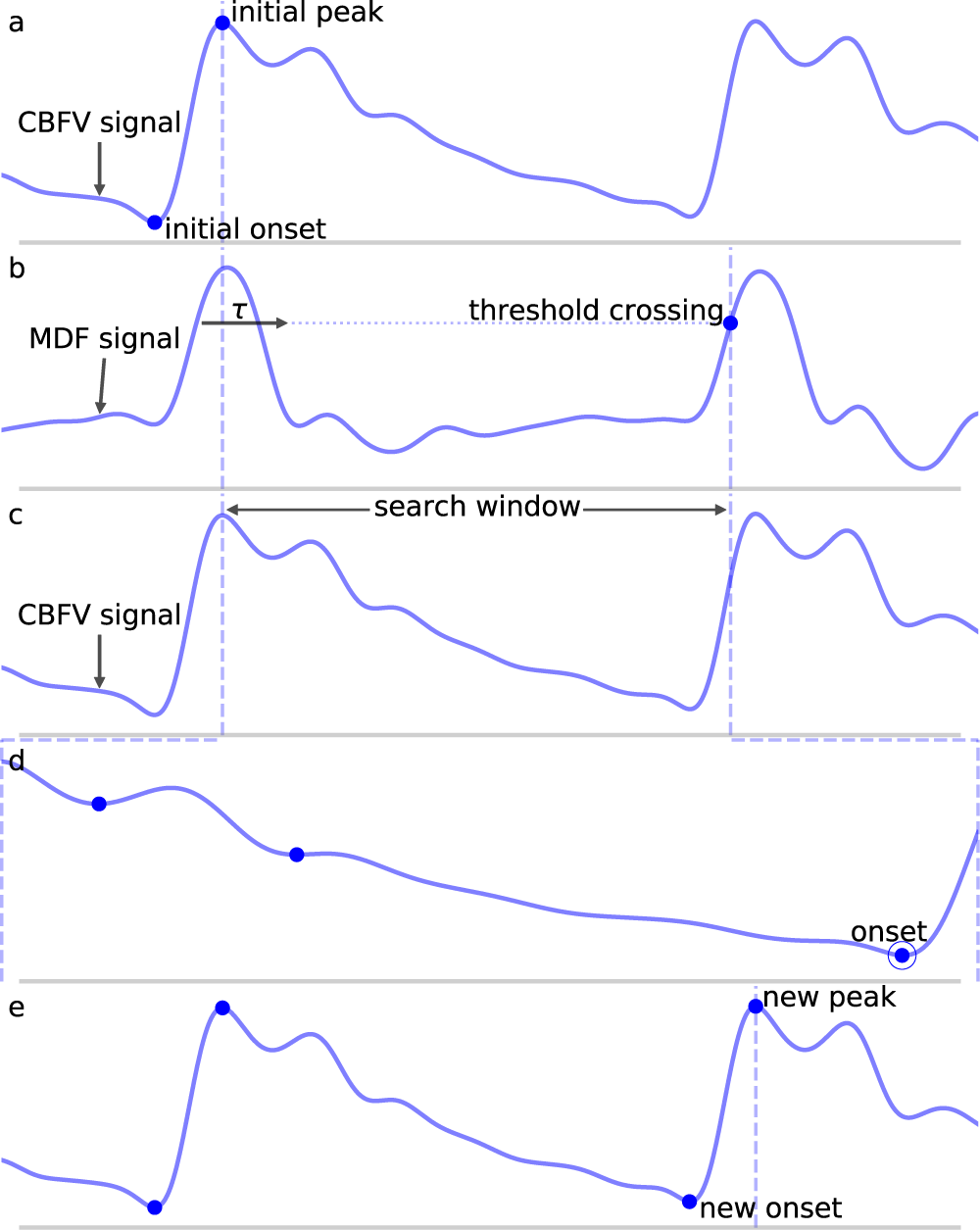
Shown are the major steps in the algorithm, from an initial seed onset to detection of the next onset. The x (time) and y (CBFV/MDF amplitude) axes have been normalized to dimensionless quantities for ease of viewing. The algorithm proceeds from onset to onset in this manner until the end of the scan is reached. Panel (a) shows the initial onset in the CBFV signal, which consists of the onset point itself and its associated peak. Panel (b) displays the MDF signal associated with this same segment of data and shows how the search window is established after enforcing the refractory period and locating the threshold crossing point. Panel (c) shows the CBFV signal with the search window overlaid. Panel (d) shows a zoom-in of the search window and the location of all valleys, with the new onset labeled. Finally, panel (e) shows the original CBFV signal with the new onset identified.

### C. Window locations

Using the MDF signal, windows in which a pulse onset are likely to occur are identified. This is accomplished by adaptively thresholding on the MDF signal. The threshold is established at 60% of the average of the preceding 20 detected peaks in the MDF signal, where a peak is defined to be the local maximum immediately following the identification of a threshold crossing point. Near the beginning of a scan prior to finding 20 peaks, the threshold value is simply calculated from all the peaks that have been found so far. The justification for using a finite number of peaks (as opposed to a single peak or all peaks) is that the properties of the signal which affect the threshold value can vary on longer time scales, so thresholds should be based on peaks in the local region preceding a particular peak. However, enough peaks should be used to guard against the possibility of overweighting transient anomalies in the signal. In practice, varying this number does not have a significant impact on the algorithm’s performance, as discussed in Section IV-B. For the initialization step, before any peaks have been found, all peaks above 2.5 times the average of the MDF signal over the first 10 seconds of data are identified, and the initial threshold is set to 60% of the median value of these peaks. We find that this process generates a reasonably accurate and robust estimate of the initial threshold.

Once a threshold value has been determined, a threshold line is propagated from the previous threshold crossing point until it crosses the MDF signal, as shown in Fig. 3(b) by the horizontal dotted line. The point where this line crosses the MDF signal is referred to as a threshold crossing point and is also labeled in Fig. 3(b). In order to avoid detecting multiple threshold crossing points associated with the same MDF peak, a refractory period *τ* is enforced immediately following the detection of a threshold crossing point, during which no new threshold crossing points can be found. The refractory period *τ* is also labeled in Fig. 3(b). The value of *τ* is taken to be 200 ms, though the precise value, once again, does not have a major impact on the algorithm’s performance (refer to Section IV-B for more details) so long as it is longer than the typical pulse upslope time but significantly shorter than an entire pulse, defined as the time between successive onsets.

A search window, shown in Fig. 3(c) between the dashed vertical lines, is then defined as the region between the location of this threshold crossing point and the peak of the last detected pulse immediately preceding each new search window. In the case of the very first onset, for which there is no preceding peak, the search window is extended to the very beginning of the signal. The peaks of each beat should occur very close to the threshold crossing point, and are determined by finding the maximum value which occurs within the refractory interval *τ*. We note here that the algorithm does not explicitly set out to accurately identify systolic peaks, and the finding of peaks is only performed in order to facilitate defining onset search windows. Thus, no claims are made regarding the accuracy of the systolic peaks.

### D. Onset identification

Once a search window has been identified, all valleys (identified by finding local minima) in the CBFV signal that occur within this window are located. The valley which occurs latest, i.e., the valley closest to the threshold crossing point, and for which the condition

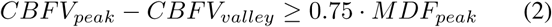

is satisfied is marked as the pulse onset. Here, *CBFV*_*peak*_ is the value of the peak for the CBFV pulse (the amplitude of the point labeled “new peak” in Fig. 3(e)), *CBFV*_*valley*_ is the candidate onset (the points marked in Fig. 3(d) with dots), and *MDF*_*peak*_ is the amplitude of the peak value of the MDF signal for this window (the value of the local maximum immediately following the threshold crossing point in Fig. 3(b)). This condition (eq. 2) ensures that the algorithm avoids mistakenly identifying valleys that appear in the upslope due to noise artifacts or pathological morphologies as onsets. The constant scale factor in front of *MDF*_*peak*_ is set to 0.75, though onset identification is fairly robust to changes in this factor (refer to Section IV-B). In theory, *CBFV*_*peak*_ – *CBFV*_*valley*_ should very nearly equal *MDF*_*peak*_ since the MDF signal measures the net change in the CBFV pulse waveform over some small time window. Thus, any factor close to but slightly less than 1 is reasonable, e.g., 0.5 – 0.95. A lower value generally provides less protection against misaligned onset detections, while too high of a value risks missing the onset altogether. The candidate onset closest to the threshold crossing point that meets the required condition is circled and labeled as “onset” in Fig. 3(d).

### E. Beat length analysis

After the signal has been scanned through in its entirety and all initial guesses at pulse onsets have been established, the final step is to deal with outlier beats. The motivation for this is based on the fact that the initial pass tends to result in two primary kinds of mistakes.

- **Long beats** typically occur when the algorithm misses a beat, which would normally result in a false negative (FN). This most commonly occurs because of some abnormality in the upslope of a beat, either because it is not very steep and fails to cross the threshold line or because it contains some sort of noise artifact that suppresses the MDF signal. An example of such a beat is shown in Fig. 4(a), where the amplitude of a beat is suppressed such that it fails to cross the threshold line.
- **Short beats** typically occur when noise in the signal results in a sharp upslope which is erroneously identified as a new beat onset resulting in a false positive (FP). The effect of this is to divide what should be a single beat into multiple short beats.

**Fig. 4.**
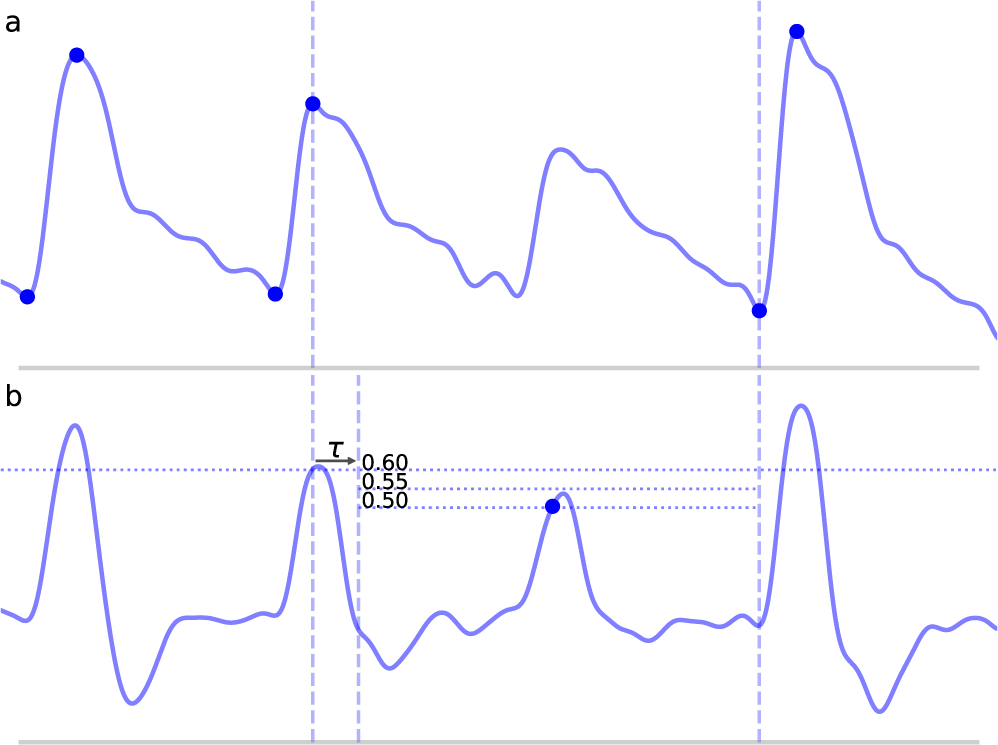
Shown above is a long beat resulting from a missed onset due to a suppressed upslope. The x (time) and y (CBFV/MDF amplitude) axes have been normalized to dimensionless quantities. The CBFV signal is displayed in panel (a), while the MDF signal is shown in panel (b). Also shown in panel (b) are the threshold values used for onset detection, which are incrementally relaxed until either an onset is found or until some minimum threshold value is reached. The search window for the new onset is indicated by the vertical dashed lines once the refractory period *τ* has been enforced.

We posit that errors of these varieties can be effectively identified by examining the distribution of beat lengths, i.e., the distance between successive onsets, resulting from the first pass of onset detection. Before proceeding however, it is worthwhile to describe a few general principles that guide a number of the choices that we make regarding the algorithm. The first principle is that we weight false negatives (failing to detect a beat onset) more severely than false positives (detecting extra beat onsets). Said another way, we believe it is better to find as many beat onsets as possible even if that means a few extra onsets are falsely identified. The motivation for this belief is influenced by the second principle, which is that the output of the algorithm should allow any downstream applications (anything using the output of the algorithm) to make the majority of the decisions about how to handle the beats once they have been segmented.

For example, in the case of short beats, these usually result from mistakenly dividing a beat into two beats, so the solution may be to just delete the onset associated with the short beat. However, this can always be done as a postprocessing step. Furthermore, such post-processing can be tailored to suit the specific needs of the downstream analysis being performed, information which is not known by the beat segmentation algorithm at runtime. Thus, as a general purpose onset detection tool, we believe it is best to be as conservative as possible, and only perform very simple post-processing steps to identify and handle the most egregious errors. This philosophy will be referenced as motivation for a number of the design decisions made in the following discussion.

#### 1) Outlier detection

In order to flag beats as outliers, we use median absolute deviation (*MAD*) due to its robustness to outliers, which was also used in [14]. *MAD* is computed according to the following:

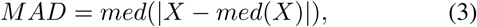

for some univariate data set *X* (the set of beat lengths in this case), composed of *N* elements *x*_1_, *x*_2_, …, *x*_*N*_, where med denotes the median value. To use *MAD* as a consistent estimator of the standard deviation, a scale factor of 1.4826 is needed, which is related to the assumed normality of the distribution after excluding outliers. Thus, the estimator *σ̂* for *X* becomes 1.4826 · *MAD*.

Finally, we define rejection criteria based on this estimator by choosing a threshold value. Setting this threshold value is a necessarily subjective choice that depends on the nature of the data and on the degree of caution deemed appropriate by the researchers. A threshold value of 3.5 has been suggested by [22]; however, a threshold value of 3.0 is also commonly suggested as a fairly conservative threshold as in [23]. In this study, we opt to use the more conservative value of 3.5 to identify short beats and a value of 3.0 to identify long beats. We believe this to be in line with the general philosophy of performing the least amount of post-processing as possible, and to deal with only the most extreme outlier cases. The choice of an asymmeteric threshold represents the goal of minimizing FN. Since long beats are generally indicative of a missed onset detection, we use a slightly less conservative threshold when identifying long beats. These choices are examined in more detail in Section IV-B. Concretely, the inclusion criteria then becomes

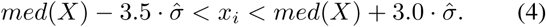

So long as *x*_*i*_ satisfies eq. 4, it is not an outlier. We can now formalize the concept of a short beat and a long beat. For a given scan for which *N* total beats have been detected with median length *l*_*med*_, for a given beat with length *l*_*i*_ and estimator *σ̂*, the beat is short if it satisfies eq. 5 and long if it satisfies eq. 6.

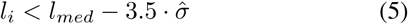

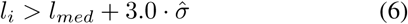

#### 2) Long beats

Long beats are dealt with first. The method for dealing with long beats is to apply the global onset detection algorithm to a local section of the signal, with a progressively lowered threshold line if needed. This process is detailed in the steps below. For each long beat detected:

i. The search window is defined from the beginning of the peak for the long beat until the next detected onset, as shown in Fig. 4(a) and denoted by the vertical dashed lines.
ii. The MDF is computed for this segment of data, as shown in Fig. 4(b).
iii. A threshold line is set at 60% of the median of all the peak heights that were located in the original global MDF signal during the first pass.
iv. After enforcing the refractory period, this threshold line is propagated until it either crosses the MDF signal or reaches the end of the data segment. If it crosses the MDF signal, then a search window is defined and new candidate onsets are located in exactly the same manner as described in the global onset detection algorithm.
v. If new onsets are found, these locations are saved and the algorithm proceeds to the next long beat in the scan or, if there are no more remaining long beats, the next step in the algorithm. However, if no new onsets are found, the threshold value is lowered by 0.05 and steps (iii) and (iv) are repeated with the lower value. The algorithm continues to lower the threshold value in this manner until either a new onset is found or some mininum threshold value is reached. A high minimum threshold value may result in more missed onsets, whereas a lower value may cause an increased number of false positives. We chose a relatively low minimum threshold value of 0.35 based on the view that false positives are preferable to false negatives (refer to Section IV-B for discussion). Fig. 4(b) shows this process of new threshold lines being drawn until the missed peak is finally detected.

#### 3) Short beats

Short beats are dealt with last. This step looks at each short beat along with each of its immediate neighbors and makes a decision as to whether the short beat should be combined with either of its neighboring beats. A number of methods can be used to determine whether a merger should occur. A simple method which we find to work effectively and which is also conservative is to simply look at whether a merger of beats would produce a new beat with a length closer to the median beat length than the original neighboring beat. If so, then the short beat is examined to determine whether it looks sufficiently “different” (as defined in the following steps) from an average beat in the scan. If both conditions are met, then a merger is performed; otherwise, no merger occurs. The process for determining this is described below.

i. Four lengths are determined: *l*_*before*_, *l*_*short*_, *l*_*after*_, and *l*_*median*_, shown in Fig. 5. These are defined as follows: *l*_*before*_ is the length of the beat occuring before the short beat, *l*_*short*_ is the length of the short beat, *l*_*after*_ is the length of the beat occuring after the short beat, and *l*_*median*_ is the median beat length of all beats that have been found in the CBFV signal for a given scan. The length of a beat is defined to be the time between successive onsets.
ii. The mean beat is calculated, shown in the subfigure of Fig. 5. This is done by first finding the median beat length. All detected beats are then either truncated or padded by repeating the last value in the beat such that all the beats have the same length, resulting in a set of length normalized vectors. The mean beat is then calculated by taking the mean of this set of vectors.
iii. The algorithm checks to see if combining the short beat with the beat occuring immediately after it, resulting in a new beath length *l*_*short*_ + *l*_*after*_, would produce a beat with a length closer to the median beat length than *l*_*after*_ alone. If it would, then one final comparison is made: the beats are combined only if the correlation distance between the short beat and the mean beat is greater than 0.2. The correlation distance is computed by truncating the longer of the two beats such that its length is equal to that of the shorter beat, and then applying eq. 7, where u and v are vectors representing each of the two beats being compared after length equalization.

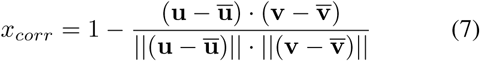 The reason for this comparison is again based on the view that false negatives are worse than false positives, and so the deletion of onsets should be done very conservatively. The motivation for enforcing this check is the idea that beats which result from noise in the signal should not look like an average beat, whereas beats that are actually just physiologically shorter will still retain the general morphological structure of a typical beat and thus should have a small correlation distance from the mean beat. If after this comparison, the beats are merged, then the algorithm moves on to the next short beat if there are any. If the beats are not merged, then the same comparison is made with the beat occuring immedaitely before the short beat. If no merge is performed there either, the algorithm proceeds to the next step.
iv. One final step occurs in which the short beat is deleted if it is determined to be extremely different morphologically than the mean beat. In this context, “extremely” different is defined quantitatively by a correlation distance greater than 0.7. The choice of this threshold and others is explored in more detail in Section IV-B.

**Fig. 5.**
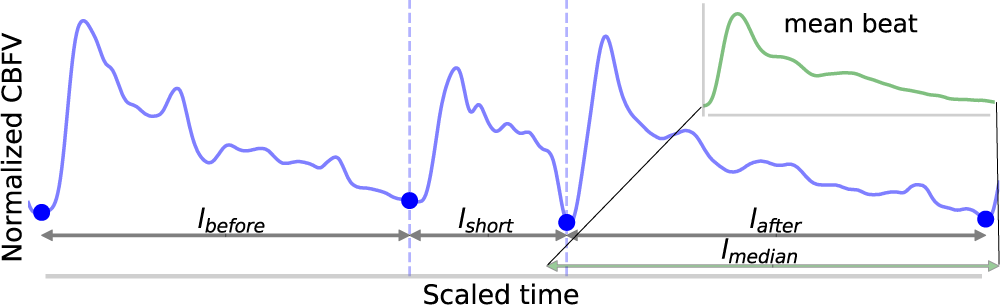
Shown above is an example of the detection of a short beat. The short beat is shown in the main figure between the dashed vertical lines between its two neighboring beats. Detected onsets are marked by blue dots. The lengths *I*_*before*_. *l*_*short*_, and *l*_*after*_ are also labeled. The mean beat calculated for this scan is also displayed in the subfigure shown in the upper right corner (not to scale), and its length *l*_*median*_ is displayed correctly to scale alongside the other distances in the main figure. The x (time) and y (CBFV/MDF amplitude) axes have been normalized to dimensionless quantities.

#### 4) Iterating

It is possible for a single pass to not find all the new beats or delete all the short beats due to the fact that as beats are added/subtracted, the statistics may change very slightly. Another way to miss a beat is due to the fact that when parsing a long beat, the algorithm will exit upon finding the first new beat. If a long beat contains more than two beats, then this means a single pass will not be enough to locate all the new beats. To deal with this, the beat length analysis is performed repeatedly until no new beats are added/subtracted during a single iteration. In very rare instances (this never occurs in the data set used in this work), it may be possible to reach an oscillating solution, so a maximum number of iterations of 10 is enforced, whereby the loop will exit regardless.

### F. Beat alignment

The final step is to fine tune the alignment of the onsets. While not strictly necessary, we find that including this step provides a significant benefit for onsets which have a large degree of misalignment relative to their annotated locations. In qualitative terms, the alignment is performed by computing a scalar value for each onset. This value represents the estimated amount by which the onset should be shifted forward or backward in time in order to better align with the actual foot position of the beat. If this estimated amount is larger than some threshold, then the shift is applied to the onset. The reason for imposing a threshold is that for large estimated shifts, we find that the shift generally moves the onset closer to its actual position. However, for small estimated shifts, corresponding to cases where the algorithm has already closely identified the actual onset, the benefit to trying to apply further realignment is questionable and in general, seems to result in slightly worse performance for very small misalignments. As a result, if the estimated amount is less than the threshold, the shift is not applied.

In order to compute the actual shift amount, the mean beat is first computed in the same way as described in Section III-E3. The beginning portion of the beat is extracted by taking the first *N* samples of the mean beat, where *N* is equal to the number of samples in the refractory period *τ*. This segment will be referred to as the mean beat upslope. The motivation for focusing on the beginning portion of the beat is the assumption that the relevant morphological feature to align on in a CBFV pulse is the initial sharp upslope. Using an extraction window equal to *τ* works well as a general rule of thumb method for capturing the initial upslope along with some of the systolic peak. Next, a set of new potential onset positions are generated by taking a window of the CBFV signal and sliding it one sample at a time, starting a time *τ* before the detected onset and ending a time *τ* after the onset. After normalizing each vector, a dot product is then calculated between each of these new shifted onset positions and the normalized mean beat upslope. The temporal separation between the detected onset and the shifted onset which maximizes the dot product is the estimated shift. If this shift is above 30 ms, then the shift is applied; otherwise, no action is performed. This threshold can be set by the researcher based on their individual misalignment tolerance.

As mentioned, this step has no effect on the number of FN or FP and serves only to better align the beats in cases of large misalignments. An exploration of alternative methods for performing this alignment could be the subject of future work.

## IV Results

### A. Performance metrics

We report on a number of metrics to evaluate the performance of the algorithm. These metrics are based on the number of true positives (TP), the number of false positives, the number of false negatives, and the temporal separation between the annotated beat and the detected beat or temporal offset error (Δ*t*). For our purposes, metrics involving true negatives (TN) are not particularly meaningful since (1) almost every point is a TN and (2) the meaning of a TN becomes ambiguous in the region immediately surrounding a true onset.

Even defining what it means to be a TP, FP, and FN can prove problematic. A common convention in the literature is to define some arbitrary threshold value Δ*t*_*max*_. If an onset is detected such that the distance between the detected onset and the nearest true onset is less than Δ*t*_*max*_ and there are no other closer detected onsets, then the detected onset is a TP. If no true onset exists within the threshold window or there is another detected onset closer to the true onset, then the detected onset is a FP. Similarly, for a true onset, if no detected onset exists within the threshold window, then this results in a FN. Where this can become problematic is in the case where an onset is detected in association with a true onset, but their temporal separation Δ*t* is greater than Δ*t*_*max*_. This situation is represented in Fig. 6(c) for Δ*t* > Δ*t*_*max*_. What is intuitively either a single mistake or potentially no mistake at all is counted as two mistakes, both a FP, because no true onset exists “near” the detected onset, and a FN, because no detected onset exists “near” the true onset. In these situations, it may be necessary to make a subjective decision about whether to classify these types of mistakes as FP or FN. This situation is not uncommon in signals with high amounts of noise or poor signal quality or for signals whose morphology diverges significantly from what would typically constitute a normal, healthy waveform. In these cases, there may be large amounts of uncertainty when manually labeling the location of the onset, but this uncertainty is ignored when enforcing a uniform and universal classification threshold.

**Fig. 6.**
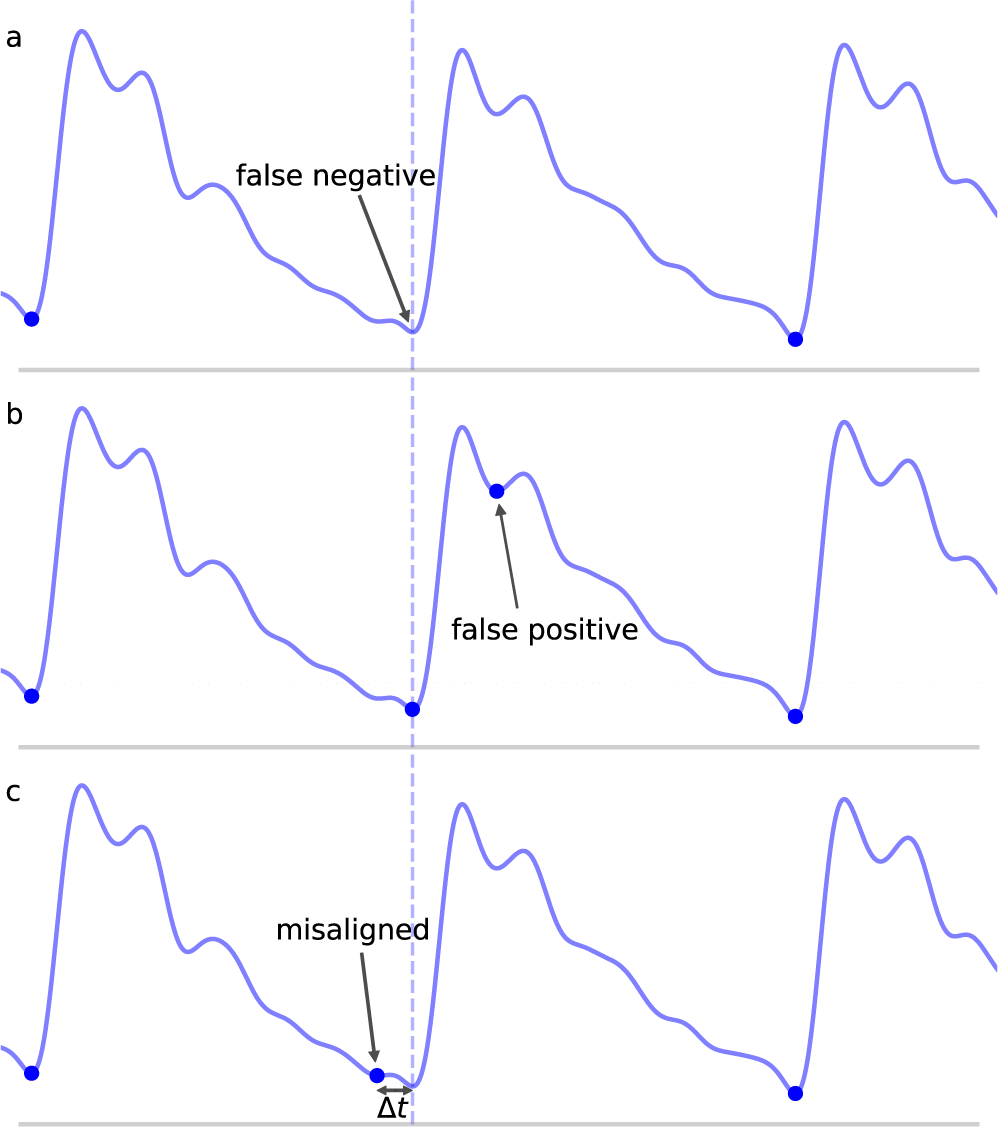
Shown are the different types of classification errors. These are shown for illustrative purposes and are not actual output of the algorithm. The dotted vertical line indicates the correct position of the beat onset. Panel (a) shows the typical situation in which a false negative arises: the algorithm fails to mark an onset for an obvious beat in between two neighboring detected beats. Panel (b) shows an example of a false positive, in which an extra onset unassociated with any true onset has been detected. Panel (c) illustrates a misaligned beat in which a true onset has been found, but its placement is offset from the true location.

For these reasons, we adopt what we believe to be a more intuitive approach to error classification in the context of beat segmentation. We avoid enforcing an arbitrary threshold value and instead focus on mapping true onsets to their associated detected onsets and then quantify the temporal offset error as a separate measure. As long as a detected onset can be associated with a true onset straightforwardly in a one-to-one manner, then the pair is labeled a TP and the temporal separation is measured. Concretely, this is done by starting with the true onsets and locating the closest detected onset, being careful to avoid double counting by ensuring that a detected onset is paired uniquely with only its closest true onset (though we note that while this situation of multiple pairings was checked for, it was never actually encountered in the results presented in this work). Any true onsets left without an associated detected onset are labeled FN. Any detected onsets left over after this process are labeled FP. A comprehensive set of unit tests were designed, and all TP pairs with large separations (> 100 ms), of which there were 44 cases, as well as all FN and FP cases, were manually inspected to confirm correct behavior of the procedure.

This classification framework naturally lends itself to three types of nominal errors: FN, FP, and a new error type we will refer to as misaligned. Examples of each of these error types are shown in Fig. 6. By classifying errors in this way, we get the following intuitive interpretations for the various types of errors: FN can be thought of as missing onsets, FP as extra onsets, and misaligned as onsets that are detected but which are offset from where they should be. The degree to which misalignment occurs is encoded by the distribution of Δ*t* for all annotated-detected onset pairs.

Algorithm performance is reported in Table I, which shows the total number of TP, FP, and FN, as well as the true positive rate (TPR) or recall defined in eq. 8, and positive predictive value (PPV) or precision defined in eq. 9. The mean temporal separation 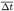 is also shown.

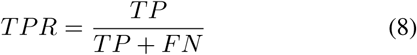

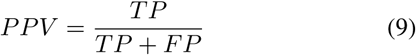

**Table I.** Shown are the relevant performance metrics of the algorithm on the entire data set.

| TP | FN | FP | TPR | PPV | $\overline{\Delta t}$ (ms) |
| --- | --- | --- | --- | --- | --- |
| 92010 | 2 | 2 | 0.99998 | 0.99998 | $6.098 \pm 4.749$ |

For the data set considered in this work, the algorithm was able to detect an onset associated with nearly every annotated beat onset (2 FN) while managing to only detect two false onsets (2 FP), again given a total of 92,012 annotated onsets, for a TPR and PPV both approaching 99.998%. This represents a significant improvement over previous work using a similar data set, which reported a TPR of 93.1% and a PPV of 93.3% [20]. It should be noted that prior work classified TP, FP, and FN using a different convention and as a result these numbers should not be compared directly; however, the gap in performance is still sizable. Using the same 30 ms value for Δ*t*_*max*_ as was used in [20], we would obtain a TPR of 99.5% using the same error classification method as used in [20].

An analysis of the failure modes of the algorithm can prove instructive into understanding the ways in which it can fail, and the combination of factors that must occur in unison to cause failures. The FP are shown in Fig. 7. These serve to illustrate the main way in which the algorithm can falsely detect beats as both cases involve a sudden upslope in the signal likely caused by noise. Normally, these would be dealt with during the beat length analysis step, as the false detections result in short beats that would normally get merged. However, in Fig. 7(a), the FP occurs in the middle of what is an unusually long beat for this scan (the local heart rate was very low relative to the overall heart rate of this scan), and so splitting it resulted in two beats that are only slightly short and thus fail to register as short beats during outlier detection. In Fig. 7(b), the beat is identified as a short beat, but because it looks morphologically similar to a typical beat, its correlation distance from the mean beat is very low and it does not meet the criteria for merging.

**Fig. 7.**
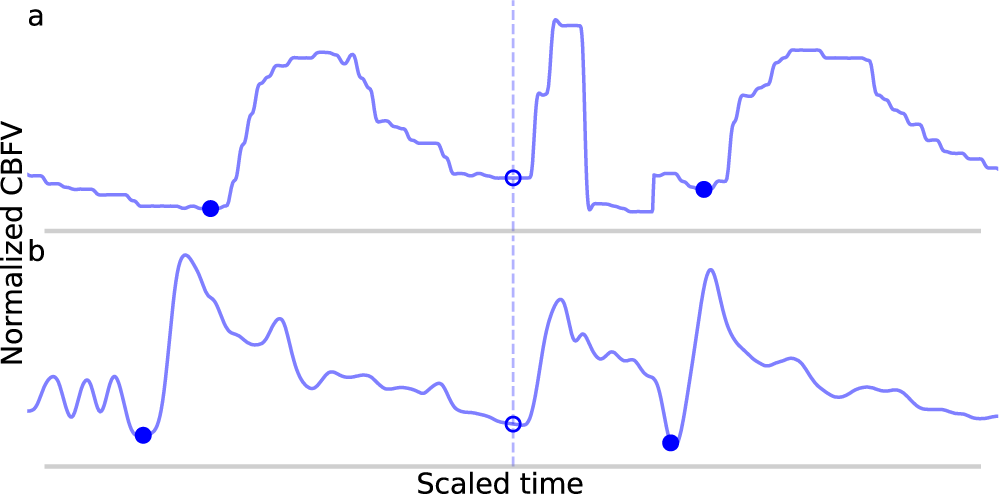
Shown in panels (a) and (b) are the false positives detected by the algorithm in the data set. The correctly identified manually annotated beat onsets are marked by filled-in blue dots. The false positives have been aligned on the dotted line and are marked by empty circles.

The FN are shown in Fig. 8. In Fig. 8(a), the onset is actually detected initially, but it just happens to be very short (perhaps indicative of a sudden, momentary increase in heartrate) and also just morphologically different enough from the mean beat that it registers for deletion. In Fig. 8(b), the onset is not detected during the initial pass due to the abnormally small difference between the beat’s systolic peak and diastolic valley, resulting in a suppressed MDF peak. The two beats which occur here locally (the missed beat and the beat immediately preceding it) are again very short, and even when combined, fail to approach the long beat threshold. It can be seen that the length of these two beats together is indeed very similar to the adjacent beats.

**Fig. 8.**
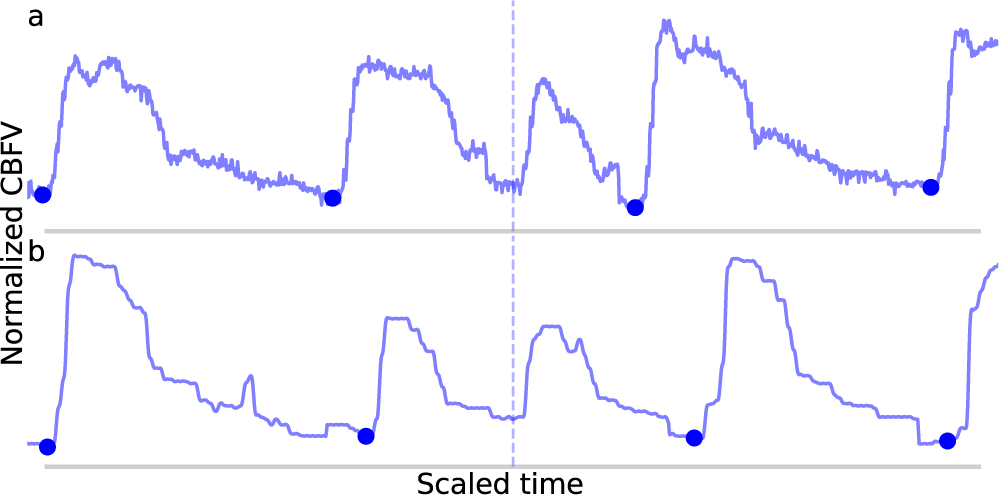
Shown in panels (a) and (b) are the false negatives detected by the algorithm in the data set. The correctly identified manually annotated beat onsets are marked by filled-in blue dots. The locations in which an annotated onset was indicated but no onset detected are marked by the dotted line.

For three out of the four errors (Fig. 7(b) and Fig. 8) described above, a case can be made that the true label is ambiguous based on manual inspection of the CBFV signal alone, and that perhaps one could reasonably conclude that these should not have been classified as errors. Indeed, based solely on the CBFV waveform, it is not clear why, according to the annotations, Fig. 7(b) is not a true onset while Fig. 8(a) and Fig. 8(b) are. However, in order to remain as objective as possible, we report only results which assume that the independently annotated labels are in fact the ground truth. Nevertheless, based on these results, it may appear that the cases in which the algorithm might fail are likely to lie at the very boundary of what would be differentible to a human expert.

Fig. 9 shows the distribution of the temporal separations between annotated-detected onset pairs. As expected, the vast majority of separations are less than 10 ms with a steep dropoff thereafter. Separations on the higher side, around 50 ms or more, tend to be the result of poor quality or pathological signals, in which there is no obvious exact point of onset, but instead a region, anywhere in which an onset could reasonably be marked based on the CBFV waveform alone. Effectively, the signal quality is such that there exists a high degree of uncertainty in the start of the beat as would be judged by an expert human. Table II gives the proportion of onset pairs whose temporal separation is less than the given threshold. Already, within 10 ms, 97.8% of onsets are detected, and by 50 ms, 99.9% are detected. This table can also be interpreted as the TPR for a given Δ*t*_*max*_ when attempting to compare results with other methods using the more conventional error classification scheme.

**Fig. 9.**
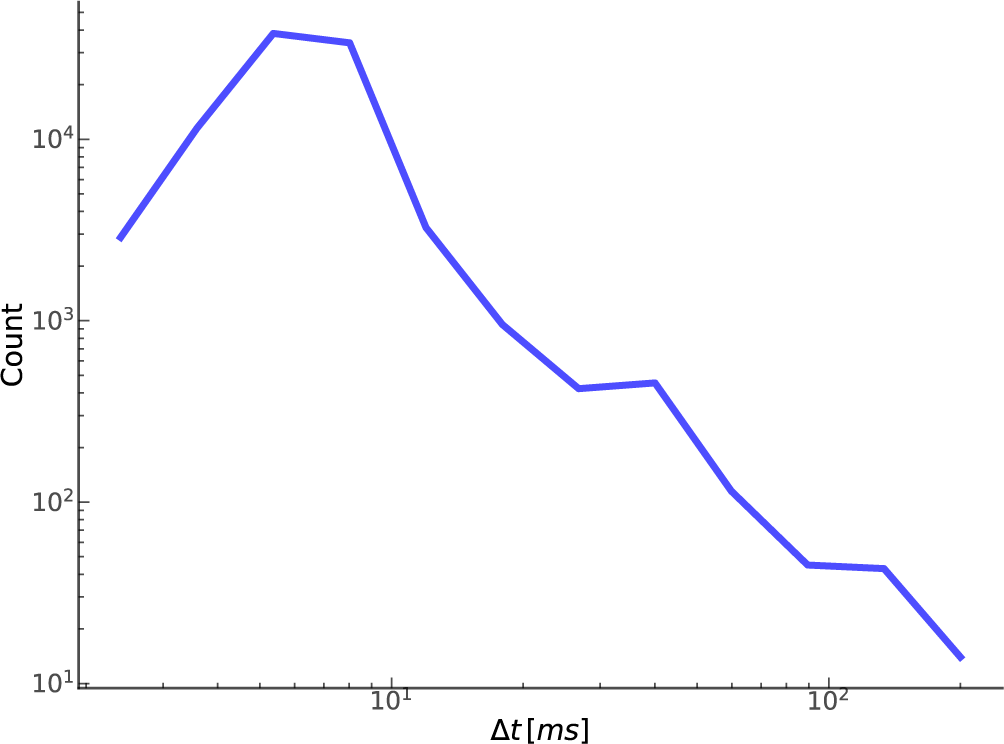
Shown here is the relationship between the number of true positives and their temporal separation.

**Table II.** Shown are the proportion of annotated-detected onset pairs detected at each threshold value. This can be thought of as a discretized representation of the cumulative distribution function.

| $\Delta t_{max}$ (ms) | Proportion |
| --- | --- |
| 5 | 0.5731 |
| 10 | 0.9778 |
| 20 | 0.9897 |
| 30 | 0.9951 |
| 50 | 0.9986 |
| 100 | 0.9995 |
| 150 | 0.9999 |

### B. Free parameters

A central claim in this paper is that the performance of the algorithm is not highly dependent on a large set of arbitrary and highly tuned free parameters. Though there are a sizable number of parameters, the algorithm appears to be very robust to changes in these parameters as long as they remain within reasonable ranges. These ranges are generally physiologically or mathematically motivated, though in some cases may need to be determined empirically. A summary of the free parameters along with a brief description of each parameter is presented in Table III, and the recommended range for each parameter as determined in this work is presented in Table IV.

**Table III.** Shown below are summary descriptions of the free parameters.

| Parameter | Description |
| --- | --- |
| Threshold normalization | Normalization factor for threshold line on MDF signal |
| Refractory period | Time to wait before a new onset can be detected |
| MDF window | Window size for computing MDF function |
| Peaks number | Number of peaks to include in MDF threshold line calculation |
| Distance threshold 1 | Correlation distance threshold for merging beats |
| Distance threshold 2 | Absolute correlation distance cutoff for deleting short beats |
| Minimum threshold norm | Minimum normalization factor to use for threshold line when processing long beats |
| Short outlier | Threshold factor to use when identifying short beats |
| Long outlier | Threshold factor to use when identifying long beats |
| Onset threshold | Minimum difference between amplitude of peak and candidate onset during onset identification |

**Table IV.** Shown below are recommended ranges for each free parameter used in the algorithm along with the actual values that were used in this work.

| Parameter | Recommended range | Value used |
| --- | --- | --- |
| Threshold normalization | [0.5, 0.65] | 0.6 |
| Refractory period (ms) | [150, 300] | 200 |
| MDF window (ms) | [125, 200] | 150 |
| Peaks number | [3, 30] | 20 |
| Distance threshold 1 | [0.05, 0.6] | 0.2 |
| Distance threshold 2 | [0.6, 0.8] | 0.7 |
| Minimum threshold norm | [0.3, 0.4] | 0.35 |
| Short outlier | [2.5, 3.5] | 3.5 |
| Long outlier | [2.5, 3.5] | 3.0 |
| Onset threshold | [0.5, 0.9] | 0.75 |

In order to demonstrate the algorithm’s robustness, we present the magnitude of the effect of varying each parameter on the performance of the algorithm. For each parameter, a range of values are tested, and the resulting performance is recorded. Due to the low number of FN and FP relative to the total number of annotated onsets, the performance will be reported in terms of the absolute number of FN and FP, as opposed to TPR and PPV, which both remain ≥ 0.999 throughout the entire range of each parameter. Due to the high dimensionality of the parameter space, an exhaustive search would be difficult, so only one parameter is varied at a time, with the rest fixed to their values given in Table IV. We feel that this is sufficient to provide a general sense of the sensitivity of the algorithm to the precise value of each parameter. The results of this procedure for nine out of the ten parameters are shown in Table V. The only parameter not listed is the peaks number, which showed no impact on the algorithm’s performance over the recommended range ([3,30]) given the data set used in this study. It should be noted that using fewer than three peaks did negatively impact the algorithm’s performance and is therefore not recommended.

**Table V.** Shown below are the values tested for each parameter and the resulting performance in terms of the number of FN and FP. The values that were actually used when reporting results are indicated by bold font.

| Threshold norm. |  |  | Refractory period |  |  | MDF window |  |  |
| --- | --- | --- | --- | --- | --- | --- | --- | --- |
| Value | FN | FP | Val. (ms) | FN | FP | Val. (ms) | FN | FP |
| 0.5 | 1 | 11 | 100 | 2 | 9 | 110 | 3 | 8 |
| 0.55 | 2 | 4 | 150 | 2 | 8 | 130 | 3 | 3 |
| **0.6** | **2** | **2** | **200** | **2** | **2** | **150** | **2** | **2** |
| 0.65 | 4 | 2 | 250 | 3 | 3 | 170 | 5 | 4 |
| 0.7 | 7 | 3 | 300 | 4 | 2 | 190 | 5 | 3 |
| Dist. thresh. 1 |  |  | Dist. thresh. 2 |  |  | Min. thresh. norm. |  |  |
| Value | FN | FP | Value | FN | FP | Value | FN | FP |
| 0.05 | 4 | 1 | 0.5 | 3 | 2 | 0.25 | 2 | 6 |
| 0.15 | 3 | 1 | 0.6 | 2 | 2 | 0.3 | 2 | 3 |
| **0.2** | **2** | **2** | **0.7** | **2** | **2** | **0.35** | **2** | **2** |
| 0.45 | 2 | 2 | 0.8 | 2 | 2 | 0.4 | 6 | 2 |
| 0.6 | 1 | 3 | 0.9 | 2 | 5 | 0.45 | 9 | 2 |
| Short outlier |  |  | Long outlier |  |  | Onset threshold |  |  |
| Value | FN | FP | Value | FN | FP | Value | FN | FP |
| 2.0 | 4 | 1 | 2.0 | 2 | 6 | 0.55 | 2 | 3 |
| 2.5 | 3 | 1 | 2.5 | 2 | 4 | 0.65 | 2 | 2 |
| 3.0 | 3 | 1 | **3.0** | **2** | **2** | **0.75** | **2** | **2** |
| **3.5** | **2** | **2** | 3.5 | 4 | 2 | 0.85 | 3 | 2 |
| 4.0 | 2 | 5 | 4.0 | 5 | 2 | 0.95 | 3 | 2 |

Based on these results, we conclude that the values of the free parameters primarily impact the edge cases, which by definition represent a small percentage of the total number of beats. This result was expected given the fact that many of the parameters are only involved during the beat length analysis and alignment steps, which by design were intended to catch outliers. Despite rather large changes in some parameter values, the performance impact was generally negligible on overall performance. The three parameters to which the algorithm appears to be most sensitive are the threshold normalization factor, the refractory period, and the MDF window. However, the fact that a single value can perform extremely well across the wide range of subjects, waveform morphologies, and signal qualities represented in this study suggests that the parameter values established in this work may generalize well despite individual variation from scan to scan or patient to patient, without the need to perform any kind of parameter tuning.

## V. Conclusion

The results presented in table I and table II are especially promising when considering the nature of the data set. The scans were performed on patients experiencing subarachnoid hemorrhage, and the overall quality of the data generally ranges from very poor to acceptable. Numerous examples of the types of noise that may commonly be present in CBFV waveforms acquired via TCD are represented in abundance as well as a host of morphological “archetypes”. Taking into account the generally poor data quality and diverse signal morphology, we feel this data set serves as a good collection of low quality signals that may be representative of the types of data collected in real world scenarios.

Nevertheless, using our error classification framework, the algorithm presented here was able to identify all but two of the 92,012 onsets present in the data set with only two false detections. Of the correctly identified onsets, over 99.5% were detected within 30 ms of the annotated onset. These performance metrics mark a significant improvement over prior attempts at CBFV pulse onset detection. Additionally, the nature of the data set gives confidence to the notion that the algorithm may generalize well in a wide range of possible scenarios or deployment environments, and that it may prove useful in any signal whose pulse onsets are marked by the presence of a sharp upslope feature, as is the case in CBFV signals. More work may be warranted to show the reliability of the algorithm across all of these potential scenarios.

The algorithm’s robustness to noise and varied waveform morphologies is encouraging for its potential use in neuro-critical care, where these conditions may be likely to occur when acquiring CBFV signals. Such conditions are also likely to reduce the effectiveness of many existing onset detection algorithms, so it is important to measure the performance of any algorithm on data which exhibits these pathological qualities. It should also be noted that, while we stress the importance of the algorithm’s performance on poor quality data, the performance is expected to improve on better quality data. Indeed, none of the errors on the test set occurred in high quality signals with normal waveform morpology.

Another important consideration is the potential application of such an algorithm in real time systems. Integral to such systems is the computational complexity of the algorithm. We find that the algorithm described in this paper implemented in Python 2.7.14 is easily able to run on the entire data set in under 8 minutes (approximately 5 ms per pulse) on a 2017 model Dell Precision 5520 laptop computer. Given this fact, it should be capable of running in real time on any relatively modern computer hardware with very little overhead. The only major caveat is that there are a number of steps that involve future knowledge of the signal, which would need to be modified to only use past knowledge. Given that these steps are only involved in the outlier detection steps, the impact of such changes are not likely to be significant. Quantitatively evaluating this performance impact could be the subject of future work.

## Acknowledgment

This material is based upon work supported by the National Science Foundation under Grant No. 1556110. Any opinions, findings, and conclusions or recommendations expressed in this material are those of the authors and do not necessarily reflect the views of the National Science Foundation.

